# DIME: Data-driven Importance MEtric Guides Localization of the Seizure Onset Zone from Intracranial EEG Data

**DOI:** 10.64898/2025.12.19.695626

**Authors:** Richard Zhang, Alan A. Díaz-Montiel, Nooshin Bahador, Milad Lankarany

## Abstract

Epilepsy affects over 50 million individuals, many of whom require surgical treatment that is dependent on accurate localization of the seizure onset zone (SOZ). Conventional SOZ biomarkers are based on strictly defined intracranial electroencephalography (iEEG) phenomena and cannot benefit from increased datasets. The scarcity of SOZ-labeled iEEG data impedes biomarker development. We introduce the Data-driven Importance MEtric (DIME) to guide SOZ localization in an interpretable pipeline that improves with ictal-labeled iEEG data. We apply DIME to an open-source dataset (n=21; 13 successful; 8 failed) for SOZ localization and surgical outcome prediction. The highest DIME-ranked electrode belonged to the clinically annotated SOZ for 69.2% of patients with successful surgery (***p <* 0.001**). DIME scores were significantly higher in SOZ electrodes than nonSOZ electrodes in both successful and failed surgeries (***p <* 0.001**), though the DIME distribution for successful cases differed from failed cases (***p* = 0.002**). DIME predicted surgical outcome with 92.3% recall and 66.7% accuracy.

## 1 Main

Epilepsy affects over 50 million individuals across the world, a condition characterized by the recurrence of unprovoked epileptic seizures [1, 2]. Among people with epilepsy, approximately 30% of them experience drug-resistant epilepsy (DRE), in which seizures persist despite two trials of appropriate anti-epileptic drugs [3, 4]. One of the most effective treatment options for DRE is resective surgery [5]. Resective surgery involves removal of the epileptogenic zone, which is defined as “the area of cortex that is necessary and sufficient for initiating seizures and whose removal (or disconnection) is necessary for complete abolition of seizures” [6]. The seizure onset zone (SOZ) is the region of the cortex where seizures initiate, and is a strong predictor of the epileptogenic zone [7, 8]. Localizing the SOZ is a critical step for successful resective surgery [5, 8, 9]. Intracranial electroencephalography (iEEG) is the gold standard for collecting data to inform localization of the SOZ, where electrodes are placed on or implanted in the brain to record electrophysiological changes [10]. However, there are significant challenges in identifying the SOZ for resective surgery. Approximately 25% of patients who undergo invasive iEEG are unable to proceed to surgery, and even among those who do complete surgery, up to 50% do not remain seizure free [8]. Thus, localization of the SOZ is a challenging but critical step for both treatment of DRE and prediction of surgical success.

Numerous tools have been developed to attempt to localize the SOZ. Due to the significant volume and complexity of iEEG data, many have aimed to leverage automated and computational analyses to help process and interpret the data. Machine learning (ML) algorithms have demonstrated significant potential in analyzing iEEG data for seizure detection, seizure phase classification, and SOZ localization [11–14]. There has been success in leveraging the network properties of the brain’s electrophysiological signals to derive features and graph representations based on functional connectivity for seizure phase classification [13]. Despite its promise, there are still several limitations in existing ML methods for processing iEEG data. Firstly, these algorithms usually utilize supervised learning methodologies, which require an abundance of accurate ground truth labels [12, 13, 15]. This fundamentally poses an issue for the task of SOZ localization, where accurate ground truth labels are difficult to obtain as even expert clinicians face challenges in identifying the SOZ [8]. Secondly, many complex ML models are black boxes that are difficult for clinicians to understand and trust, posing an obstacle to clinical application [11].

In addition to data processing tools, several biomarkers have been proposed to assist in SOZ localization and/or prediction of surgical success. Neural fragility is a metric that models iEEG data as a dynamical system, with electrodes as nodes, and identifies which nodes are more susceptible to creating imbalances in connectivity [9]. Neural fragility is associated with nodes in the clinically annotated SOZ, and the metric is able to predict surgical success with 76% accuracy [9]. Grinenko et al. (2017) [16] extracted spectrogram features during the pre-ictal-to-ictal transition as a biomarker or “fingerprint” of the epileptogenic zone. The methodology attempted to identify electrodes within the epileptogenic zone, with 90% of its predictions matching the clinically annotated zone. Ictal chirps, another spectrogram feature consisting of low voltage narrow-band ictal fast-activity, have also been proposed as EEG markers of the epileptogenic zone [17]. Others have focused on utilizing distinct epileptic patterns in iEEG data, such as high frequency oscillations [18].

Although these metrics show promise in assisting SOZ localization, there are still needs for improvements in performance in order for them to be applied to clinical practice. Additionally, these metrics are based on static definitions or mathematical formulations. Their performance in SOZ localization will not improve unless their underlying formulations are changed, which is nontrivial to implement. Implementing error-correction algorithms for these static formulations to reduce false or missed detection of biomarkers from noise can be challenging as well. These methods, unlike machine learning approaches, cannot benefit from increases in data as their performance is independent from the dataset.

In this paper, we propose the Data-driven Importance MEtric (DIME) which associates with the SOZ and demonstrates potential for improvement with increases in ictal-labeled iEEG training data. We leverage machine learning explainability tools to develop an interpretable pipeline that can ultimately assist in SOZ localization and prediction of resective surgery success without being reliant on SOZ labeled data. Specifically, we process iEEG data into graph representations [13] that are used to pretrain a graph neural network for seizure detection (not SOZ localization), which we then use as input to an explainability algorithm that ultimately computes DIME importance scores. For our experiments, we first optimize parameters for DIME based on a general and customizable pipeline, which also serves to showcase the potential for DIME to improve and learn from training data. We then use an optimized version of DIME and assess the metric’s association with the SOZ, its ability to predict surgical success, and its relation to relevant clinical markers.

## 2 Results

SOZ localization is critical for successful resective surgery, but there are few clinically validated biomarkers for the SOZ [9]. Moreover, existing biomarkers have fixed definitions that are unable to benefit from the wealth of iEEG data and machine learning advancements accumulated over the past several years.

We propose DIME as a metric of the SOZ that can learn from ictal-labeled iEEG data. We first describe the generalizable DIME generation pipeline, and then we evaluate how it can be adjusted to optimize final computed DIME scores. Once optimized, we apply the metric to various tasks relevant to SOZ localization. For all experiments, we utilize an N=21 patient open-source multi-centre iEEG dataset from OpenNeuro [19] (see Methods for more detail).

### 2.1 Description of DIME (Data-driven Importance MEtric)

DIME is based on the idea that brain regions within the SOZ are the first to display seizure-like electrophysiological activity. Manual localization of the SOZ by clinicians often involves thorough examination of iEEG data to find electrode channels that are the first to develop abnormal or seizure activity at the time of seizure onset [20]. Our ultimate goal is to localize the SOZ by identifying which electrodes represent initial seizure activity.

An explainer function E*_t_*: (G*_t_*, f*_θ_*, Y*_t_*) → (A_*t*_^′^, N_*t*_^′^) generates scores indicating the most important features in A and N from an individual graph representation G*_t_* given a pre-trained graph neural network f*_θ_* and its decision Y*_t_*. **Step 3.** An aggregator function A *_t, t_*_=*k*_: {A_*t*_^′^ }*^T^* → L aggregates graph network importance scores across iEEG channels and across different time windows, from k to T, generating the DIME scores for each iEEG channel, L. **Interpretability.** Top: Graph network importance scores (A_*t*_^′^) rep-resent which electrode functional connections contributed most to a given decision Y. They can be thresholded to identify important electrode subgraphs involving connectivity with SOZ electrodes. Bottom: Similarly, electrode features (N_*t*_^′^), representing iEEG frequency bands, can be thresholded to identify important bands for spectrogram analysis. The information can be correlated with other spectrogram biomarkers such as ictal chirp [17]. **Final output.** DIME scores can identify which electrodes are likely to reside in the SOZ. They can also be processed to predict whether resective surgery will succeed or fail given a set of clinically annotated SOZ labels. When combined with the interpretability component, the DIME pipeline can also assist in spectrogram analysis.

We conjecture that the predictions produced by graph neural networks trained for seizure detection are attributable to certain key nodes in the input graph that correspond to electrodes exhibiting salient seizure activity. If we examine input graph representations and model predictions near the time of seizure onset, we hypothesize that model predictions are heavily influenced by electrode channels with initial seizure activity or, in other words, channels within the SOZ. Moreover, we expect that in the training process for seizure detection, these models will become more fine tuned at recognizing, processing, and understanding the complex and subtle patterns within the iEEG data associated with seizure onset. As models are trained on more ictal-labeled iEEG data, they become better differentiators of ictal and non-ictal iEEG data by learning motifs associated with seizure vs. nonseizure activity, increasing their classification performance. Thus, we hypothesize that as model performance for seizure detection increases as a result of more seizure detection training data, these models will become better at identifying the SOZ electrodes that display initial seizure activity near the time of seizure onset. In theory, DIME can *learn* from and improve at SOZ localization with more seizure detection data.

The major task then is to uncover which electrode channels our models are attributing their predictions to. To do so, we employ explainability algorithms which aim to identify the most *important* input nodes for our graph neural networks. These *importance* scores from model predictions near the time of seizure onset are ultimately used to derive DIME, which can then be used as a marker for the SOZ.

To summarize broadly, DIME is a biomarker of the SOZ that can improve with training data. DIME is derived from explainability scores that identify which electrode channels are most important for a graph neural network to make predictions for seizure detection. Though trained on seizure detection data, DIME is used for SOZ localization and resective surgery outcome prediction. Notably, DIME is not necessarily computed from any one algorithm, but is rather computed from a generalizable pipeline with components that can be adjusted. The DIME pipeline is depicted in Figure 1, and can be formulated as follows:

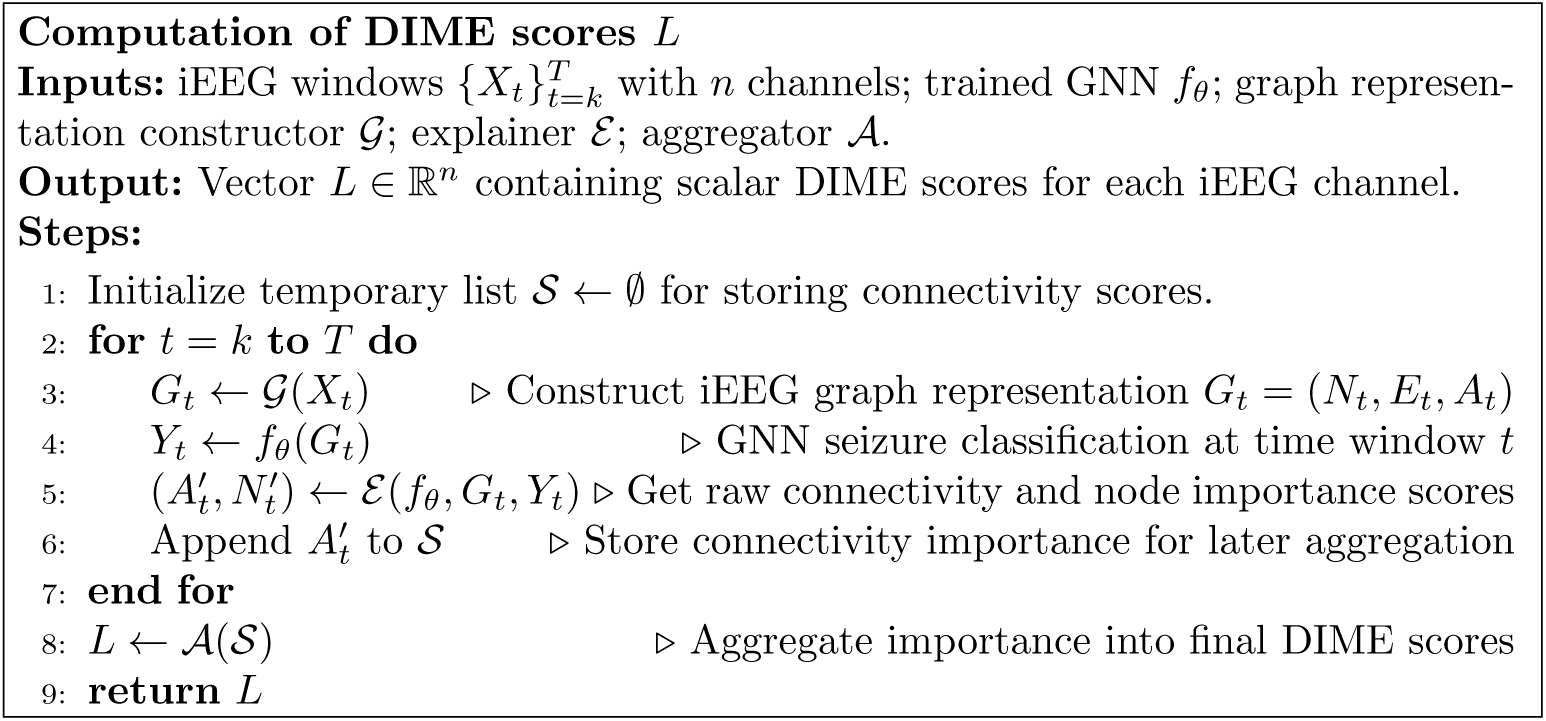

**Fig. 1:**
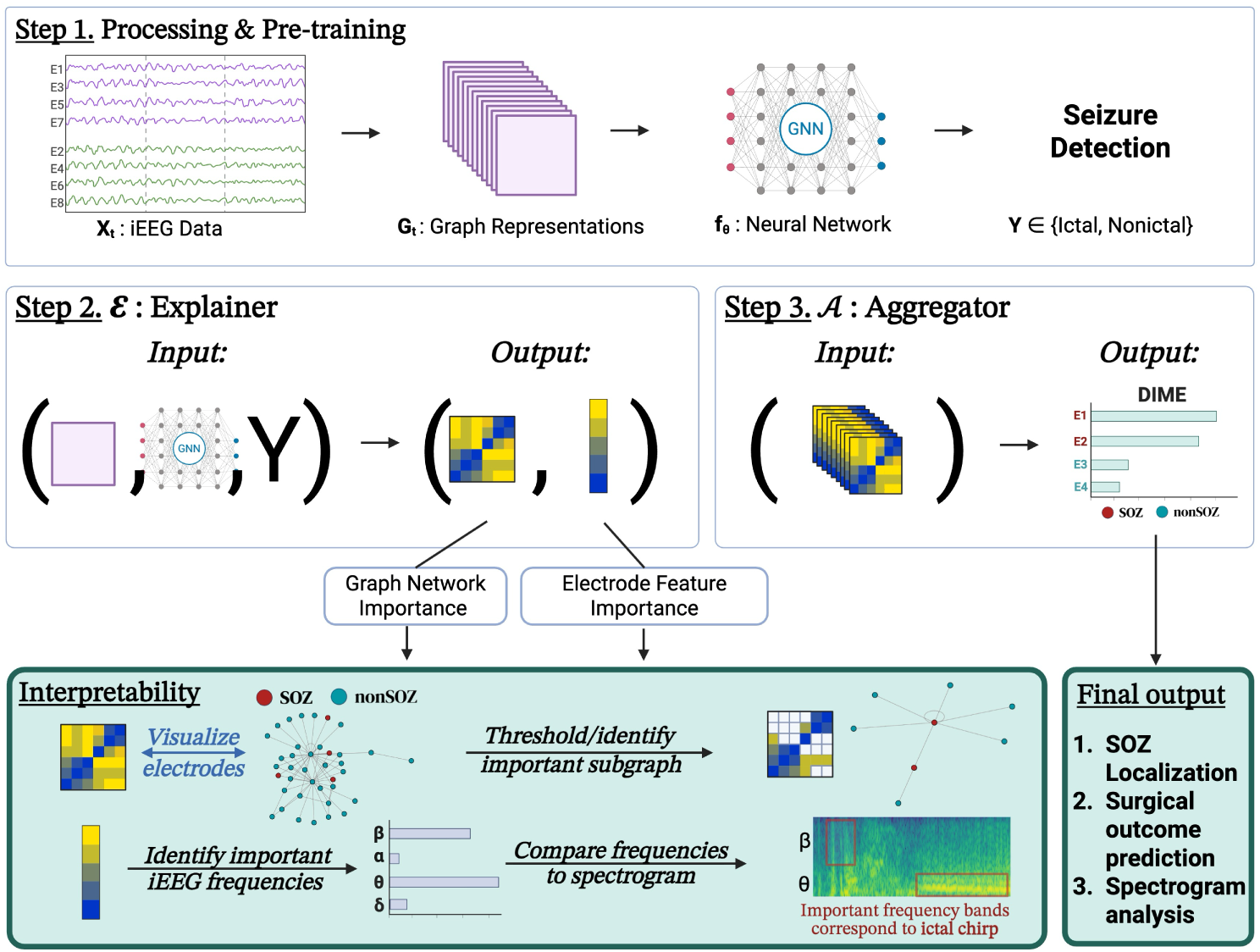
Summary of DIME pipeline. **Step 1.** Multi-channel time series iEEG data X is split into time windows t that are each transformed into graph representations G*_t_* = (N, E, A), where N ∈ R*^n×FN^* denotes F*_N_* node features (e.g., spectral power by frequency band) extracted from n iEEG channels, E ∈ R*^n×n×FE^* denotes F*_e_* edge features, and A ∈ R*^n×n^* is an adjacency matrix derived from functional connectivity measures such as correlation or coherence between channels. G*_t_* is inputted into a graph neural network f*_θ_* with learnable parameters θ and pretrained to classify the seizure label Y ∈ {0, 1}, indicating the presence or absence of seizure activity. **Step 2.**

For a comprehensive description of the methodology used to compute DIME scores, see Methods.

### 2.2 DIME learns from training data and can be optimized

A key advantage of the DIME pipeline is that it is highly flexible and trainable, rather than being a rigidly defined biomarker. We can modify any of the following processes to adjust our DIME scores: G(X*_t_*) to generate the graph representation, f*_θ_*(G*_t_*) to process the graph representation, E(f*_θ_*(G*_t_*)) to explain the neural network, or A({E*_t_*}*^T^*) to aggregate the scores. Even the underlying data X*_t_* can be augmented to improve the performance of the G(X*_t_*) and f*_θ_*(G*_t_*) operations. Functionally, any of the above modifications act with the shared similar purpose of altering the DIME score that is computed. We aim to assess how changes in DIME scores are associated with differences in SOZ localization accuracy. Without loss of generality, we examine how changes in f*_θ_*(G*_t_*) affect SOZ localization accuracy.

As demonstrated in Table 1, increases in GNN seizure detection accuracy correlate with increases in SOZ localization accuracy. Moreover, the statistical significance of the pipeline at distinguishing SOZ vs. nonSOZ nodes becomes stronger as p-values decrease. This suggests that models performing better at seizure detection generate more powerful embeddings and raw importance scores that lead to improvements in SOZ localization. The optimal f*_θ_*(G*_t_*) was the GAT-ECC GNN architecture, which yielded the greatest seizure detection and SOZ localization performance. For the subsequent sections of this paper, we will use this architecture to probe the capabilities of the DIME pipeline.

**Table 1:** The performance of the DIME pipeline when integrated with different potential GNN architectures.

| Architecture in DIME | Seizure Detection Acc | 1° SOZ Acc | Importance $p$-value |
| --- | --- | --- | --- |
| Null Hypothesis | 75% | 19.4% | 1 |
| GAT | 86.6% | 15.4% | 0.7 |
| ECC | 97.5% | 61.5% | < 0.001 |
| GAT-ECC | 98.2% | 69.2% | < 0.001 |

Possible $f_\theta(G_t)$ architectures include a graph attention layer (GAT) [21], an edge-conditioned convolutional layer (ECC) [22], or both (GAT-ECC). Seizure detection accuracy refers to the GNN performance on classifying ictal vs. nonictal, while 1° SOZ accuracy denotes the accuracy with which the top-ranked electrode by DIME score falls within the clinically annotated SOZ. The importance $p$ -value refers to the probability of obtaining a given SOZ accuracy if the DIME pipeline was making random predictions. The null hypothesis architecture is a dummy baseline whose seizure detection and SOZ accuracy performance is based on random predictions.

It is important to note that although this section’s focus is to examine how the core GNN architecture in DIME can be optimized, the other parameters for DIME have also been selected to optimize for seizure detection accuracy, based on the results of previous work [13]. The exact parameters of the DIME pipeline used in this paper, including the underlying graph representation features, GNN hidden layers, and GNN hyperparameters, are described in Methods.

### 2.3 DIME is associated with the SOZ

Using the optimized DIME pipeline with the ECC-GAT model, the highest DIME-ranked electrode belonged to the clinically annotated SOZ 69.2% (95CI 42.4%-87.32%, Wilson score interval) of the time (9/13 subjects), which was significantly higher than the null distribution of there existing 19.4% (160/826) clinically annotated SOZ electrodes in the dataset (p < 0.001, Pearson’s χ^2^; p < 0.001, Fisher’s exact test). There are clear trends that electrodes belonging to the clinically annotated SOZ are often ranked higher than those that are not (Figure 2 right).

**Fig. 2:**
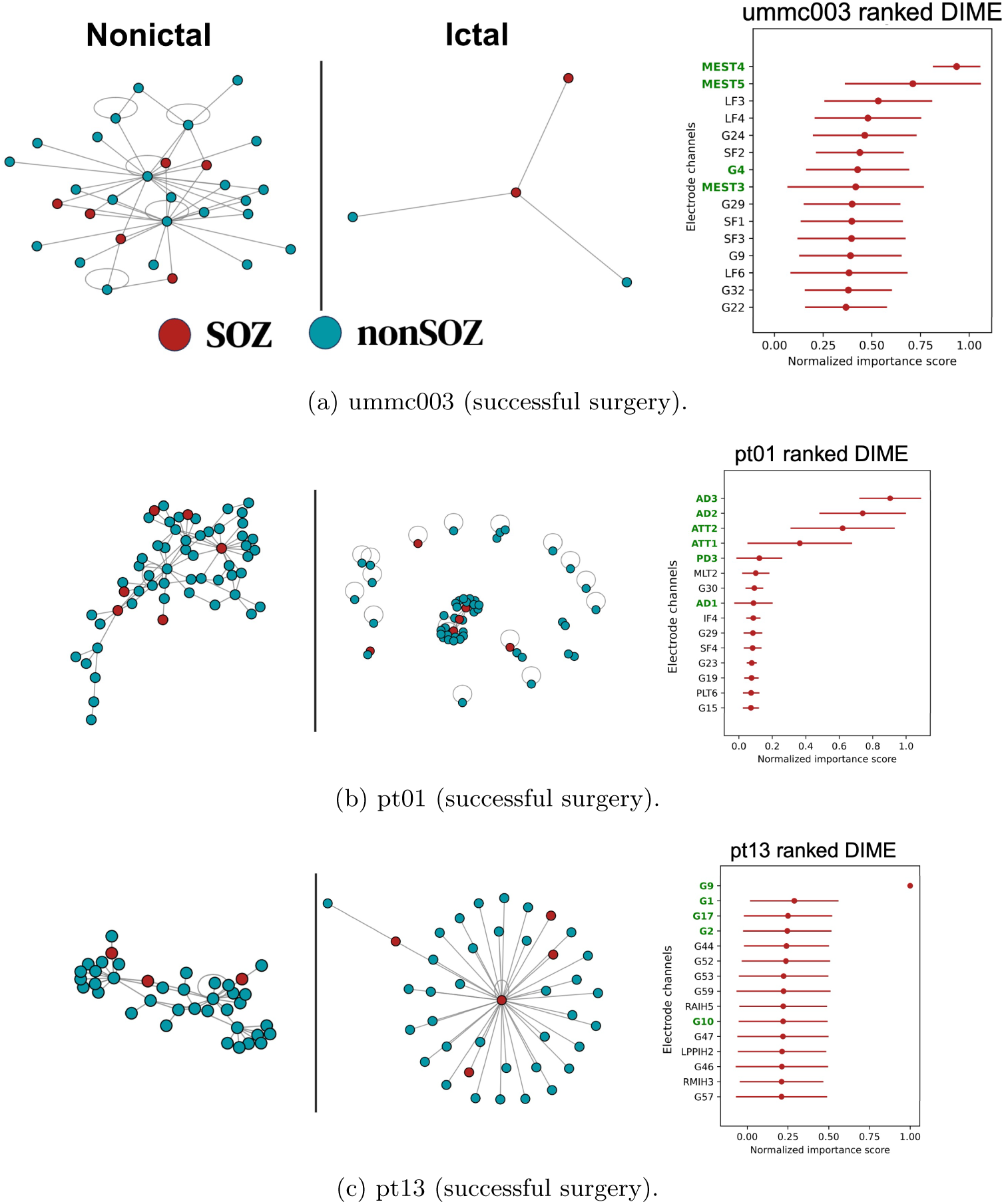
Left: Examples of nonictal (left of bar) and ictal (right of bar) subgraphs from GNNExplainer after thresholding normalized importance above 0.95. This change in connectivity, involving a transition to strong connections with SOZ nodes, is consistent for all successful surgical subjects; a representative sample is shown. Right: Top 15 electrodes ranked by DIME scores, with clinically labeled SOZ highlighted in green. Horizontal bars represent standard deviation of the non-aggregated importance scores.

We next assessed how the top 3 DIME-ranked electrodes were distributed within and outside the clinically annotated SOZ (Figure 3). Among successful surgeries (Figure 3a), the median (Q1, Q3) DIME score of SOZ electrodes was 0.843 (0.345, 1.000), whereas for nonSOZ electrodes it was 0.370 (0.161, 0.801), which was significantly different (p < 0.0001, Mann-Whitney U Test). Among failed surgeries (Figure 3b), there was a median (Q1, Q3) DIME score of 0.757 (0.330, 1.000) for SOZ electrodes and 0.512 (0.143, 0.910) for nonSOZ electrodes, which was also a significant difference (p < 0.0001).

**Fig. 3:**
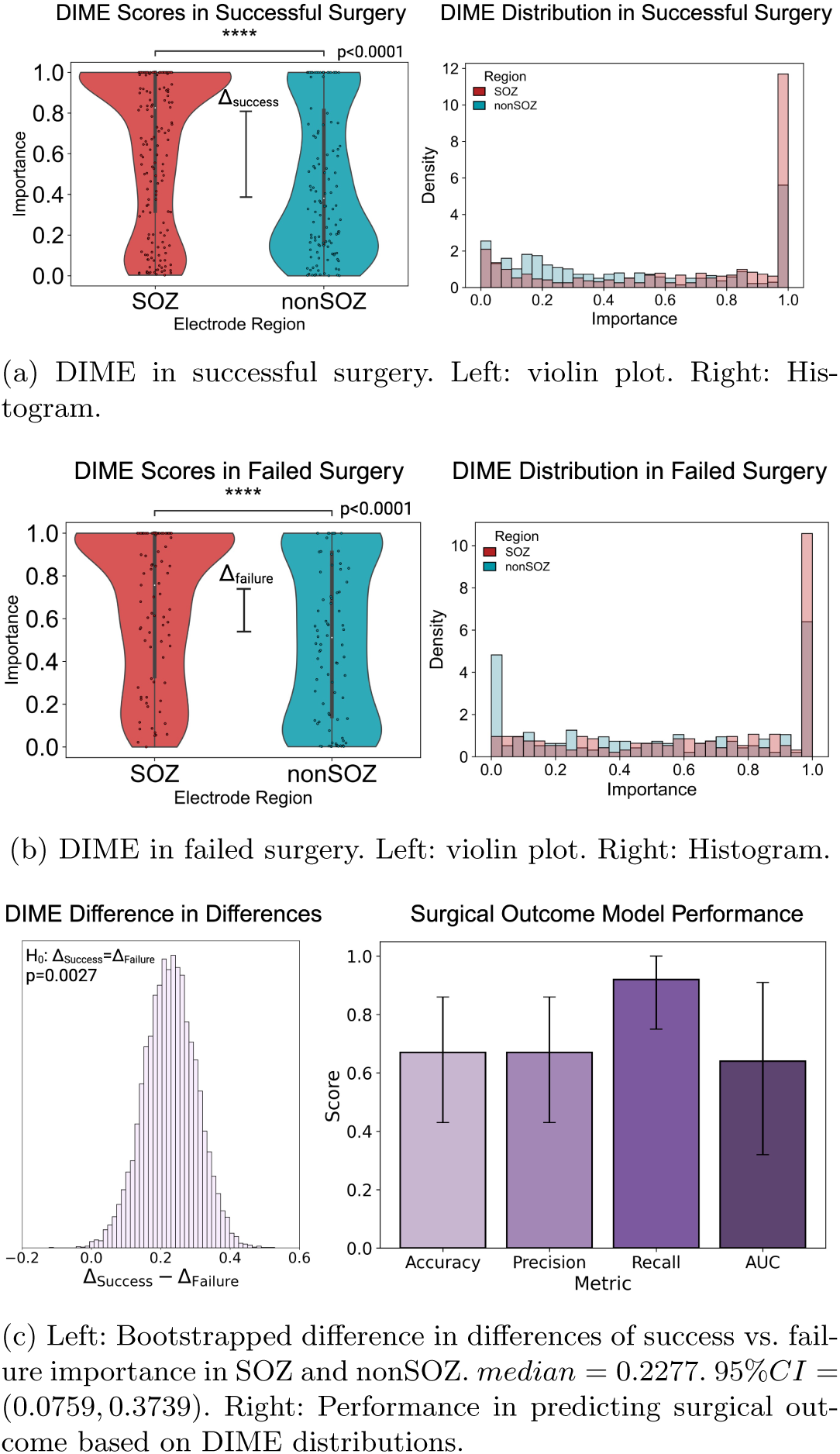
DIME score distributions. Mann-Whitney U tests were used for statistical comparisons.

When visualizing networks of the most important electrode connectivity subgraphs (interpretability component in Figure 1), a recurrent motif persists throughout multiple different subjects (Figure 2 left). In the transition from nonictal to ictal windows, there is consistently a change in connectivity involving strong connections to particular electrodes within the clinically annotated SOZ. Though the exact community structure may differ, the ictal subgraphs of all subjects include this motif of strong SOZ connectivity.

### 2.4 DIME is associated with clinical markers relevant to SOZ resection

We first assessed how DIME scores were related to surgical outcome. We examined the difference in differences of DIME scores between SOZ labels and surgical outcomes. Let us define Δ as the difference in median DIME scores for clinically annotated SOZ vs. nonSOZ electrodes, as depicted in Figures 3a and 3b on the left. Δ*_success_* was 0.473 while Δ*_failure_* was 0.245, yielding an observed difference in differences (Δ*_Success_* − Δ*_F ailure_*) of 0.228.

To determine the statistical significance of these raw values, we conducted non-parametric bootstrap resampling [23] (10,000 iterations with replacement) on the distribution of Δ*_Success_*− Δ*_F ailure_*, yielding the histogram in Figure 3c left. The difference in differences was significantly different than the null distribution (H_0_: Δ*_Success_* = Δ*_F ailure_* ⇐⇒ Δ*_Success_* − Δ*_F ailure_* = 0) with p = 0.0027. In other words, Δ*_success_* was significantly different than Δ*_failure_*; the distribution of DIME scores among the clinically annotated SOZ and nonSOZ was more segregated in successful surgeries than failed ones. This is also qualitatively observable in Figures 3a and 3b.

We ran a support vector machine model to predict surgical outcome of success or failure based on distributions of DIME scores in the clinically annotated SOZ and nonSOZ (Figure 3c right). The performance was evaluated using leave one out cross validation. Confidence intervals of performance metrics were estimated by bootstrapping the model’s set of predictions, using a similar methodology as described in Tsamardinos et al. (2018) [24]. The model notably had a high recall (95%CI) at 0.923 (0.750-1.000). Accuracy (95%CI) was 0.667 (0.429-0.857). Precision (95%CI) was 0.667 (0.438-0.875). Area under the receiver operating curve (95%CI) was 0.644 (0.316-0.914).

Lastly, we assessed the feasibility of the DIME pipeline to identify important frequency bands in the context of spectrogram analysis (interpretability component in Figure 1). Our aim was not to perform a rigorous population-level analysis, but rather to conduct a qualitative exploratory assessment to establish face validity of the pipeline methodology. Frequency band importance scores were extracted for representative subjects pt01 and pt13, who both had successful surgery (Figures 4a and 4b). Spectrograms from their electrodes were also computed, with annotations added to highlight features consistent with other validated iEEG biomarkers corresponding to the SOZ and/or epileptogenic zone (Figures 4c to 4f).

**Fig. 4:**
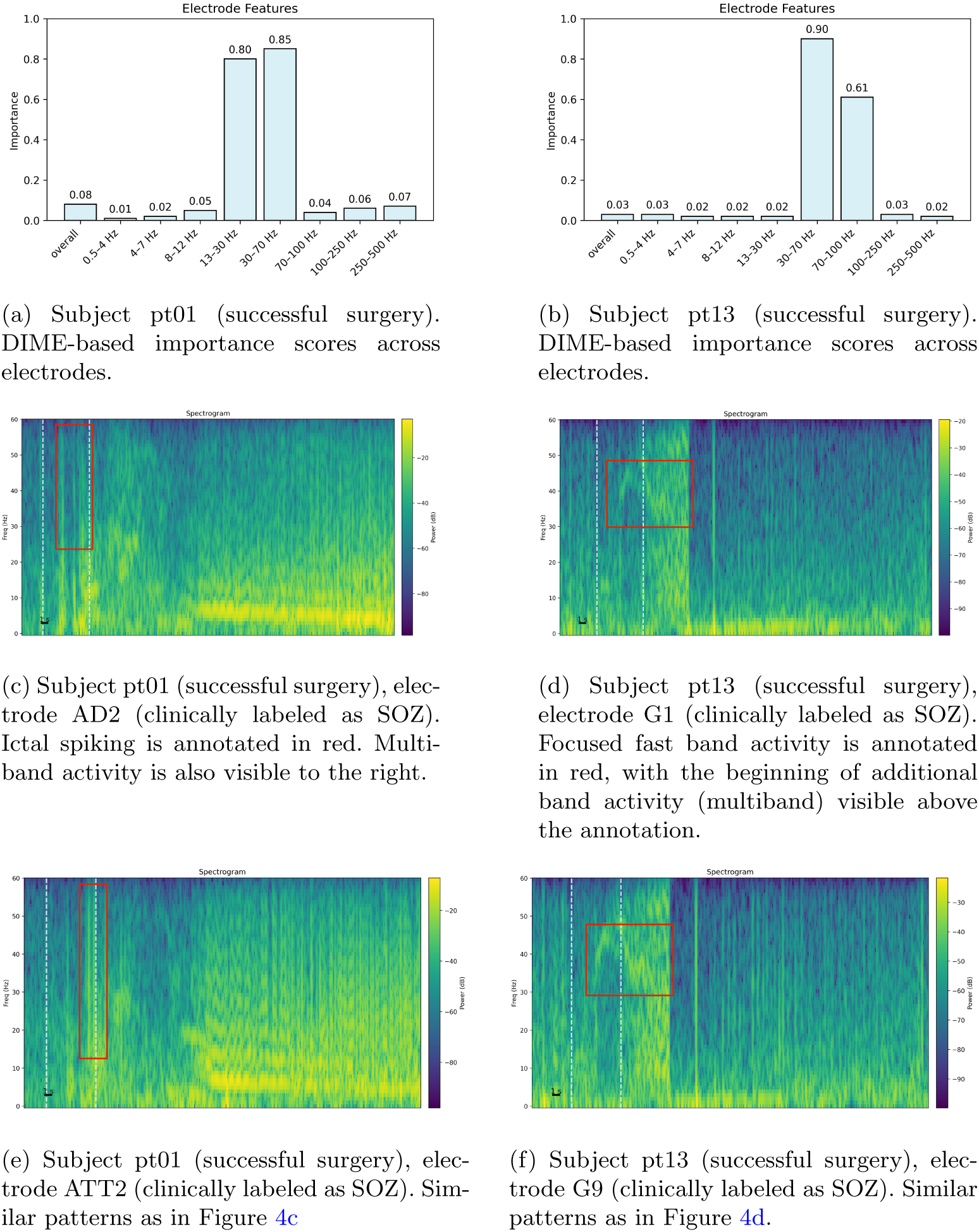
Electrode frequency importance maps and spectrograms illustrating associations with features of other iEEG biomarkers, such as the fingerprint of the epileptogenic zone [16] and/or ictal chirp [17]. The blue dotted vertical lines enclose the window used to extract the electrode frequency importance maps.

## 3 Discussion

We introduced DIME as a metric with significant associations with the SOZ that leverages GNN explainability to be able to improve with ictal-labeled training data. DIME is not statically defined based on any epileptic and/or iEEG phenomena, but is rather generated through a series of generalizable components that can learn from iEEG training data. We trained and applied one possible DIME configuration to a n=21 open-source patient dataset, where the pipeline demonstrated statistically significant performance in SOZ vs. nonSOZ localization, discrimination of success vs. failed surgery through score distributions, and prediction of surgical outcome. The important frequency bands generated by the DIME pipeline correlated with features consistent with other biomarkers, further validating its use as a marker for the SOZ. Overall, DIME appears strongly associated with the SOZ, and, crucially, has compelling potential in its ability to improve with further training data and/or augmentation of each component of its pipeline.

DIME localized the SOZ and discriminated successful vs. failed surgeries with high statistical significance. One of the challenges of SOZ localization is the sparsity of available training and validation data. This not only makes training models challenging, but it adds an extra layer of difficulty in validating their performance. With small sample sizes, there is a greater risk of model bias, random sampling error, and result imprecision, all of which pose challenges to trusting the results from experimentation. We attempted to overcome this by relying on statistical approaches, such as hypothesis testing and bootstrapping, to generate p-values and probabilistic distributions for our results to lend them greater confidence. The statistical significance of the performance by DIME is one of its strengths. Despite only having a 21 subject dataset, statistical testing, with adjustments to accommodate small samples, enables greater faith in results. We attempted to be as conservative as possible in our statistical testing, such as by prioritizing nonparametric tests that do not require data normality assumptions, to avoid type 1 (false positive) errors. Thus, DIME’s capacity for SOZ localization, specifically of its highest ranked electrode, its separation of SOZ vs. nonSOZ, and its discrimination of successful vs. failed surgeries can be taken with reasonable credibility given all of their very low p-values (p ≤ 0.002). Generally, this suggests that DIME is strongly associated with clinical interpretations of the SOZ, since our SOZ labels were all based on clinical labels. There is also potential for DIME scores to be associated with the true SOZ or epileptogenic zone, rather than only clinical interpretations, given its ability to segregate successful vs. failed surgeries. Its discriminatory ability may be a result from its capability to represent discrepancies between clinical labels of the SOZ and true labels. Taken together, these results suggest that DIME can be a strong biomarker for the SOZ.

One of DIME’s strongest results was its ability to predict surgical success with relatively high recall (0.92). This is on par with other state of the art metrics, such as neural fragility or interictal epileptogenic networks, which have recalls between 0.70 to 0.90 and AUCs around 0.80 to 0.80 [9, 25]. Recall, also referred to as sensitivity in binary classifications or true positive rate, measures the ability of the model to detect positive cases, which in this use case is a successful surgery. A high recall performance enables models to identify/predict negative cases, or surgical failures, with high confidence. A model prediction of a surgical failure is less likely to be a false negative (i.e., a potential successful surgery) due to the model’s strong ability to classify successful surgeries as such (true positive rate). For high-risk medical procedures such as surgical resection in epilepsy, high recall of the successful outcome is important to identify potential surgical failures. A DIME-based prediction of a failed surgery can indicate to clinicians, with strong confidence given the high recall performance, to be more precautious. It can suggest that the location of the SOZ may need to be reconsidered. However, it is important to note that it may also imply reasons other than SOZ localization why the surgery is likely to fail, such as perhaps indicating a difficult surgical resection process. Nonetheless, this information can be useful for clinicians to integrate into their holistic approach for treatment. DIME’s overall accuracy for surgical outcome prediction was reasonable at 0.67. This is comparable to clinician performance and other state of the art methods (e.g., neural fragility, the interictal epileptogenic network) that range between 0.5-0.75 [9, 25, 26].

DIME’s most remarkable characteristic is its trainability and modifiability. In terms of trainability, in this paper we demonstrated a trend that increases in seizure detection performance correlated with SOZ localization performance. The theoretical rationale behind this finding could be that a model that is better trained on seizure detection is stronger at linking general seizure features with input graph representations. Input electrodes exhibiting early seizure activity are more likely to be both identified and prioritized by the model such that when its decisions are explained, these electrodes will correspond to higher importance scores. By better identifying electrodes with early seizure activity, the DIME pipeline performs stronger at identifying the SOZ electrodes which participate in initial seizure activity. In fact, the primary method of SOZ localization by expert clinicians also similarly involves identifying electrodes with particular seizure onset patterns [27–29], which parallels the DIME pipeline. Thus, by training better seizure detection models with more ictal-labeled iEEG data, we can yield DIME pipelines with stronger SOZ localization performance. The usage of ictal-labeled data, rather than SOZ-labeled data which can be much more challenging to obtain as even clinical labels of the SOZ may be inaccurate due to the difficulty of localizing the SOZ, supports a pipeline that can benefit from the wealth of iEEG data available through global research efforts [19, 30, 31]. Overall, one of the main advantages of DIME as a biomarker is its ability to benefit from iEEG data with relatively convenient labels. The goal of this paper was not to identify the optimal DIME configuration, but rather introduce the general architecture of DIME, outline its general components which may be modified, and suggest proof of concept for acting as a biomarker of the SOZ. If even a baseline configuration can be a strong biomarker of the SOZ that is statistically significant and relatively comparable to other state of the art methods, a further trained, modified, and optimized configuration would have substantial potential. In terms of modifiability, there are many opportunities to alter the pipeline, such as by improving the graph representation and GNN architecture, changing the explainability algorithm that is used, and/or redefining the aggregation function. Current and future innovations for GNNs [32–34] and explainability algorithms [35–37] can enhance the DIME pipeline.

The frequency band importance scores from the DIME pipeline also correlated well with other biomarkers on spectrograms (Figure 4). For both the AD2 and ATT2 electrodes with pt01 (Figures 4c and 4e), a pattern consistent with the epileptogenic fingerprint described in Grinenko et al. (2017) [16] is seen in the spectrogram: spiking near the time of ictal onset, followed by multiband fast activity with concurrent lower frequency suppression(right of the spectrogram figures). The frequency band importance scores for pt01, which highlighted the 13-70Hz range, matched the frequency of the ictal onset spiking, which was the only fingerprint feature that overlapped with window used for the DIME pipeline. The multiband fast activity and concurrent suppression was outside of the DIME window, which may explain why frequencies corresponding to those features (0-12Hz), were attributed with low importance scores. Similarly, for pt13 in both of their shown electrodes (Figures 4d and 4f) there is a visible ictal chirp pattern comprised of low voltage fast-activity near the time of ictal onset that also overlaps with the window for DIME computation. The DIME pipeline signaled high importance for the frequencies between 30-100Hz, which matches the location of the ictal chirp. Both the epileptogenic fingerprint and ictal chirp have been validated as novel strong biomarkers for the SOZ in the current state of epilepsy research [27, 38, 39]. Thus, the ability for the DIME pipeline to correlate with these metrics lends credibility to the approach. Moreover, given this ability to correlate with other biomarkers, DIME has the potential to discover new spectrogram biomarkers and phenomena that relate to the SOZ.

It is important to place DIME in the context of other biomarkers that have been proposed for the SOZ. Traditionally, clinicians have assessed particular iEEG patterns, including low-voltage fast activity, diffuse electrodecremental events, preictal/ictal rhythmic spiking, and bursts of spiking or wave activity [27]. One major challenge of manual clinical inspection is the abundance of data needing to be processed and analyzed, impairing clinical efficiency. Thus, there have been many attempts at developing new biomarkers leveraging automated data processing. Some promising markers include high frequency oscillations [40–44] and spiking [45, 46] during interictal recordings. Aside from concerns about these markers’ sensitivity, precision, and reliability-which has been heavily debated [47–50]-these markers are unquestionably imperfect, and enhancing them would be challenging as they are restricted by their underlying definitions. This difficulty exists for biomarkers of ictal recordings as well, including connectivity-based approaches such as correlation or coherence [51–53], epileptogenic fingerprints [16], epileptogenicity [54]. Machine learning approaches to SOZ localization and biomarker generation are either supervised or unsupervised. Supervised approaches usually involve training on SOZ-labeled data [41, 55, 56], which can be challenging of labels for the SOZ are scarce, subjective, or uncertain. Unsupervised methods do not rely on labels, but often rely on identification of other validated biomarkers such as HFOs or connectivities [56]. Additionally, unsupervised methods can be black boxes where interpretability and hence clinical trust and adoption may be difficult. Overall, various biomarkers exist for the SOZ, though they are all imperfect in terms of SOZ localization performance, are statically defined so they cannot benefit from additional training data and/or remain uninterpretable. To the best of our knowledge, DIME is the sole biomarker that leverages explainability, is flexibly defined, and can benefit from ictal-labeled training data. All current biomarkers demonstrate associations with the SOZ, but the challenge is that none are strongly associated enough to be able to identify the SOZ at performances that are capable of being integrated into real-world clinical care, which is why very few of these many biomarkers are used in clinical practice [9, 27]. The SOZ localization and surgical outcome performance of DIME is similar to these other biomarkers, but the major contribution of this work is not DIME’s performance alone but rather the idea that DIME can improve with further data, whereas the other biomarkers cannot.

Ultimately, DIME has the potential to be modified, trained, and improved to a performance that may be compatible with real clinical care. By highlighting electrodes with high likelihood of residing in the SOZ, it can make clinical workflows for SOZ localization more efficient and accurate. Additionally, through predicting surgical success or failure, it can flag cases to support clinical safety. It is important to note that DIME, or any other SOZ biomarker, is not intended to completely automate and replace the existing clinical workflow of SOZ localization. Rather, it is intended as a tool to supplement existing clinicians to augment their decision-making. This is one of the reasons why explainability is critical for a SOZ biomarker; interpretability is necessary for any biomarker to be both trusted and effectively wielded by clinicians.

It is important to note that this work has limitations. One of the biggest challenges is the low sample size of the dataset used. Although there exists a wealth of iEEG seizure data, there are many logistical and technological barriers in accessing it such as patient confidentiality and ethics approval. That is why this work leveraged an open-source dataset. However, still, there are a limited number of open-source iEEG datasets that also include clinical labels for the SOZ. Although the DIME pipeline itself does not require SOZ labels, which is one of its strengths, labels were required in this work for the sake of validating the pipeline. This limited the number of datasets and subjects able to be used, leading to smaller sample sizes. As aforementioned, we attempted to manage these sample sizes by relying on conservative statistical testing, using simplistic modeling with data balancing to prevent overfitting, and running a leave one out cross validation protocol to improve generalization. However, certain issues are difficult to avoid with low sample sizes, such as the fact that epilepsy is a heterogeneous disease with varying severity and mechanisms. Different epileptic or seizure etiologies may be over or underrepresented in the sample, potentially influencing experimentation results and external validity. Another limitation was the ambiguity associated with the SOZ. We assessed DIME performance based on the clinically annotated SOZ, but these may not necessarily be correct for both successful and failed surgeries. However, we supposed that there was a greater likelihood of the clinical SOZ labels being correct in successful surgeries, which is why we examined SOZ localization performance mainly in the context of successful cases. We attempted to compensate for this by assessing not only SOZ localization but also surgical outcome prediction. The task of surgical outcome prediction treats SOZ labels as an input feature rather than an outcome to validate on, preventing false results. However, there are still issues with this as there are many factors affecting surgical success (as defined in the dataset) that may be beyond accurately localizing the SOZ, such as surgical technique, disease severity, comorbidities, and follow-up time. These factors were not able to be controlled for, either because information was unavailable or because the small dataset would not allow for more complex modeling to adjust for extra co-variates.

Future work can aim to address these limitations by applying the DIME pipeline to datasets with greater sample sizes, potentially iEEG data that is locally collected if external datasets are insufficient. The problem of limited data may become less of an issue as concerted efforts to streamline the sharing of clinical iEEG data for research is growing rapidly. Additionally, the DIME pipeline can be expanded on by attempting to modify its various components and/or train it on new data to optimize its configuration.

In all, DIME is a SOZ biomarker that is unique from others in its ability to improve with further modification and data. It represents an expansion of the traditional understanding of biomarkers from strictly defined phenomena to flexible and fluid constructs.

## 4 Methods

The entire end-to-end pipeline is summarized in Figure 1.

### 4.1 Dataset

We utilized a publicly available multi-center dataset from OpenNeuro (Accession Number: ds003029) [19]. Data collection at each participating site was approved by the respective institutional review board (IRB): University of Maryland School of Medicine IRB; University of Miami Human Subject Research Office—Medical Sciences IRB; National Institutes of Health IRB; Johns Hopkins IRB; and Cleveland Clinic IRB. Informed consent was obtained from all participants at each site, and the acquisition of research data did not interfere with patients’ clinical care or objectives.

The dataset comprises intracranial EEG (iEEG) and scalp EEG recordings from 100 individuals across five epilepsy centers in the United States, all of whom were candidates for resective surgery. However, a substantial portion of the dataset was unavailable because one research center did not properly de-identify and release its data. The OpenNeuro repository currently contains 35 accessible patient entries. Of these, 12 were excluded from our analysis due to inconsistencies in clinical annotations—such as missing seizure onset or offset event markers—which are essential for our supervised GNN model framework. Additionally, two subjects (ummc001 and ummc007) were excluded because they did not undergo resective surgery. Consequently, our final dataset included 21 subjects.

All included subjects were implanted with electrocorticography (ECoG) electrodes. Surgical outcomes were categorized as successful if the patient remained seizure-free following surgery, and unsuccessful if seizure activity persisted. The duration of post-surgical follow-up varied between patients, ranging from 1 year to 7 years. Further details and clinical features of this data are described in past work on the open source dataset [9, 13].

### 4.2 Graph representation and neural network pipeline

This section refers to step 1 in Figure 1. We first pretrain a graph neural network on iEEG data to detect seizures (i.e., classify ictal vs. nonictal). In order to benefit from the functional connectivity and network information from iEEG data, we leverage the graph representation and neural network pipeline from [13]. Briefly, the original iEEG data is processed and partitioned into one second windows, where various functional connectivity metrics, including coherence and correlation, can be combined with electrode features, including energy and energy by frequency band, to derive a graph representation. Thus, a graph representation consists of connectivity metrics for its edge features and adjacency matrix as well as electrode metrics for its node features. Based on the results of [13], for this work we use an optimized graph representation setup comprised of energy by frequency band node features, correlation and coherence by frequency band edge features, and coherence as the adjacency matrix metric. The node and edge features are computed from a 1s window size, while the adjacency matrix uses a 20s window size. The window step size was 0.125s.

These graph representations are used to train a graph neural network for seizure detection. Depending on experimental setup, the neural network may have an edge-conditioned convolutional layer, a graph attention layer, or both [21, 22]. These layers were previously found to have the greatest performance for seizure detection in a graph representation pipeline [13].

Further details on the initial iEEG processing, graph representation, and graph neural network pipeline can be found in previous work explaining the pipeline [13].

### 4.3 Graph neural network explainability

This section refers to step 2 in Figure 1. Once we have a trained graph neural network for seizure detection, we use it as input to a single instance neural network explainer algorithm. Although there exist many possible explainer algorithms with their own unique advantages and disadvantages, we employed GNNExplainer, which is a well-established algorithm that has shown strong performance [57]. GNNExplainer is a model-agnostic interpretability method designed to explain predictions made by GNNs. It identifies the most influential subgraph structure and node features that contribute to a particular prediction. Specifically, it optimizes a mask over both edges and features to maximize the mutual information between the masked subgraph’s representation and the model’s output. In doing so, GNNExplainer produces an interpretable explanation showing which connections and attributes were most important for the model’s decision.

The key outputs of this component are importance scores for the input graph’s adjacency matrix (A*_t_^′^*) and node (N*_t_^′^*) features. Depending on the purpose of the DIME pipeline, one or both outputs can be used. For instance, if hypothetically the sole task was to localize the SOZ without further interpretability or analysis, an explainer algorithm that only produces adjacency matrix scores without node scores could be viable. In this work’s case with GNNExplainer, both A*_t_^′^* and N*_t_ ^′^* are computed, enabling a full exploration of SOZ localization, surgical outcome prediction, and spectrogram frequency analysis, as discussed in the following sections of the methodology.

We used GNNExplainer as a straightforward and robust algorithm for the sake of demonstrating viability of the DIME metric, but further work may aim to optimize the explainability module of the pipeline using other methods of graph node and/or adjacency explainability [14, 35, 36, 58].

### 4.4 Aggregation function

This section refers to step 3 in Figure 1. The ultimate purpose of this module is to combine all of the raw importance scores (A*_t_^′^*, N*_t_ ^′^*) generated by the explainer module E for each graph representation window G*_t_* into a single usable metric for each electrode.

This may involve aggregation across the temporal dimension t, and/or aggregation across spatial dimensions (e.g., collapsing A_*t*_^′^∈ R*^n×n^* into R*^n^*such that n electrodes each have one scalar score) In this work, we used the following function for A:

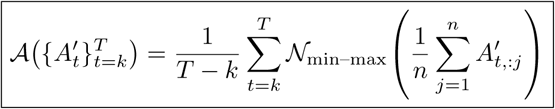

where:

A_*t*_^′^ ∈ R*^n×n^* is the symmetric connectivity importance matrix at time window t, 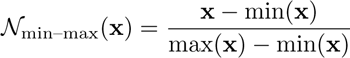 rescales the vector **x** elementwise to the range [0, 1].

The final output is L ∈ R*^n^*, which represents the final DIME vector containing singular values for all n electrodes. This aggregation function, which utilized mean temporal and spatial aggregation as well as min-max normalization, was both an intuitive way to collapse matrix/vector dimensions as well as efficient to compute. Thus, this formula was used for all experiments to serve as a simple aggregator with the purpose of demonstrating feasibility of the DIME pipeline.

A critical parameter of all aggregation functions is the selection of which graph representation windows (corresponding to particular times t) to use. Based on empirical exploration, we found that DIME scores correlated best with the clinically annotated SOZ when t corresponded to the first 25 windows after ictal onset, which is what we used for all reported experiments. In other words, we set k =time of ictal onset and T = k + 25. This aligns with what would theoretically be expected to be the most feature rich time segment for the SOZ, as clinicians also often focus on iEEG data around the time of ictal onset during their attempts at SOZ localization.

Similar to the explainability module, the aggregation function is flexible to modification as long as the fundamental purpose of summarizing raw scores into a singular metric is upheld. Again, future works may aim to optimize methods for aggregation.

### 4.5 Interpretability

This section refers to the interpretability component of the DIME pipeline in Figure 1. For visualization of an important subgraph, the key step is to threshold which connectivities to display in A_*t*_^′^. Raw importance values range from 0-1. The exact threshold value likely varies depending on the underlying data and can be found through empirical testing and exploration. Our subgraph visualizations had the best balance of connectivity information without excessive cluttering at a threshold of 0.95, which is what was used in Figure 2.

For electrode feature importance scores, the outputs of the explainer algorithm directly correspond to the importance of different frequency bands as long as the input graph representation used energy by frequency band as node features. Frequency band importance information can be used to assist or guide spectrogram analysis. For spectrogram visualization, iEEG data was processed using Fourier transform-based methods [59].

### 4.6 Application of DIME

This section refers to the final output of the DIME pipeline in Figure 1. DIME can be used to identify electrodes with high probabilities of residing in/near the SOZ by ranking electrodes based on DIME score. Higher ranked electrodes, such as the top-1 or top-3, can have high probabilities of residing within the SOZ.

The DIME distribution among clinically annotated SOZ and nonSOZ labels can also be used to segregate resective surgery outcomes. The statistical significance of the difference between the distributions for successful vs. failed surgery can be assessed with bootstrapping.

To attempt to predict surgical outcomes, we used a support vector classifier, with a Gaussian kernel to handle nonlinearity. Class weights were automatically balanced, scaling each class inversely proportional to its frequency in the training data, to address potential Y class imbalance in the training data. The input to this model consisted of a 4 dimensional vector comprised of the DIME median among SOZ electrodes, DIME IQR among SOZ electrodes, DIME median among nonSOZ electrodes, and DIME IQR among nonSOZ electrodes. From a feature engineering perspective, the dimensionality of the input aimed to be minimized to prevent overfitting to the n=21 sample size. Future works with greater dataset sizes may explore enriching the input vector with more features and/or using more complex predictive models. We employed leave one out cross validation where from the dataset of n=21 subjects, the model was trained on a subset of n=20 subjects and attempted to predict the outcome of the excluded validation subject. This process was repeated to generate out-of-set validations predictions on all n=21 subjects, and these predictions were used to evaluate performance in terms of accuracy, precision, recall, and AUC.

Finally, the results of DIME and its associated interpretability components can be compared with other biomarkers, such as ictal chirp [17], to improve spectrogram analysis.

### 4.7 Statistical analysis

Descriptive statistics were presented with means, medians, standard deviations, interquartile ranges, and confidence intervals. Wilson score intervals were used to estimate confidence intervals for proportions due to the small sample sizes [60]. For two-sample comparisons, nonparametric testing was preferred whenever possible in order to conservatively account for the possibility of non-normal distributions in the data. With continuous variables, we used Mann-Whitney U tests which are non-parametric and do not require an assumption of data normality. For comparisons of categorical variables and/or proportions, we used Pearson’s χ^2^ tests. For small sample sizes (expected counts ≤ 5 in a contingency table), Fisher’s exact test was also employed. We used a significance threshold α of 0.05.

To estimate confidence intervals and hypothesis testing values for parameters without clear distributions, bootstrap resampling with replacement was conducted [23, 61]. We also estimated confidence intervals of regression performance by bootstrapping model predictions using the methodology in Tsamardinos et al. (2018) [24], but adapting it to our models by using only a singular hyperparameter configuration and skipping all steps for hyperparameter tuning/selection.

## 5 Data availability

All data produced are available online at the OpenNeuro Epilepsy-iEEG-Multicenter-Dataset with accession number ds003029 [19], accessible through https://openneuro.org/datasets/ds003029/versions/1.0.7.

## 6 Code availability

The entire DIME pipeline-from generating graph representations with raw iEEG data to computing DIME scores and visualizing important subgraphs or frequency bands-has been containerized into a Python package, available at https://github.com/Richqrd/bgreg-main-public. To use the repository, sample scripts are located in the ‘projects’ directory and the Python modules are located under ‘bgreg’. Input data can be inserted under ‘data files’, enabling dataset customization.

## Acknowledgments

We thank the epilepsy monitoring unit at the Toronto Western Hospital in Toronto, Canada for sharing their experience in dealing with epilepsy patients and in explaining the challenges in bridging data scientists to epilepsy care.

## 7 Author information

### 7.1 Contributions

R.Z., A.A.D.M., and M.L. were involved in study conceptualization; R.Z. designed and conducted the methodology, data analysis, and results visualization; M.L. verified the study concept; A.A.D.M. assisted in data interpretation; N.B. assisted in data processing and visualization; R.Z. and M.L. drafted the original manuscript; R.Z. and M.L. were involved in manuscript review and editing; M.L. provided funding support and project supervision.

## 8 Ethics declarations

### 8.1 Competing interests

The authors declare no competing interests.

## Notes

https://openneuro.org/datasets/ds003029/versions/1.0.7

https://github.com/Richqrd/bgreg-main-public

### Competing Interest Statement

The authors have declared no competing interest.

